# Phenotypic Analysis of GGDEF/EAL Domain Protein Functions in Phytopathogenic *Pantoea ananatis*

**DOI:** 10.64898/2026.05.12.724576

**Authors:** Okhee Choi, Yeyeong Lee, Byeongsam Kang, Yongsang Lee, Jinwoo Kim

**Affiliations:** Institute of Smart Space Agriculture, Gyeongsang National University, Jinju 52828, Republic of Korea; Department of Plant Medicine, Gyeongsang National University, Jinju 52828, Republic of Korea

**Keywords:** cyclic-di-GMP, diguanylate cyclase, *Pantoea ananatis*, pellicle, phosphodiesterase, swimming

## Abstract

Cyclic diguanosine monophosphate (c-di-GMP) is a ubiquitous bacterial second messenger that regulates diverse cellular processes, including colony morphology, motility, biofilm formation, and virulence. It is synthesized by diguanylate cyclases (DGCs) containing the GGDEF domain and degraded by phosphodiesterases (PDEs) containing the EAL domain. However, studies on the genetic and physiological characteristics of c-di-GMP metabolism in *Pantoea ananatis* are lacking. In this study, we identified 26 predicted c-di-GMP metabolism-related genes in the *P. ananatis* PA13 genome: 9 encode GGDEF-only domain proteins, 5 encode dual GGDEF/EAL domain proteins, and 12 encode EAL-only domain proteins. We constructed overexpression strains and mutants of 26 DGC- and PDE-encoding genes, and then assessed their Congo Red binding, mucoid and rugose phenotypes, pellicle formation, and swimming motility. We identified 14 of 26 DGC and PDE proteins that affect phenotype changes. Among the 26 DGC- and PDE-overexpressing strains, 13 exhibited the phenotypic changes described above, with some showing alterations in multiple phenotypes simultaneously. Notably, overexpression of *dgcM* induced changes across all phenotypes. Among the 26 DGC and PDE mutants, the *pdeC* mutant increased pellicle formation and Congo red binding, the *pdeM* mutant reduced the mucoid phenotype, and the *pdeS* mutant, which shows high similarity to *ydiV*, an anti-FlhD factor, increased swimming motility. Overexpression strains and mutants of 14 DGC and PDE proteins that exhibited phenotypic changes had higher intracellular c-di-GMP levels than the wild type. This study provides important insight into the role of the c-di-GMP network in the plant pathogen *P. ananatis*.

**IMPORTANCE:** *Pantoea ananatis* is a versatile bacterium that causes significant diseases in various economically important plants. To survive and infect hosts, bacteria use a key signaling molecule called c-di-GMP to switch between swimming freely and forming protective communities known as biofilms. Despite its importance, the specific genes governing this signaling network in *P. ananatis* remained unknown. In this study, we systematically identified and characterized 26 genes responsible for regulating c-di-GMP levels in *P. ananatis* PA13. By analyzing mutants and overexpressing these genes, we pinpointed 14 critical factors that control essential behaviors such as motility, pellicle formation, and colony appearance. Notably, we discovered specific genes, such as *dgcM* and *pdeS*, that act as master regulators of these traits. This comprehensive functional map of the c-di-GMP network provides essential insights into how this pathogen adapts to its environment, offering potential targets to control plant infections.

## INTRODUCTION

Cyclic diguanylate (c-di-GMP), which acts as a nucleotide second messenger in many bacterial species, regulates a wide range of functions, including cell differentiation, biofilm formation and sessile lifestyle, motility, and virulence (1,2). Cyclic-di-GMP is synthesized by diguanylate cyclases (DGCs) and degraded by phosphodiesterases (PDEs).

DGCs, containing the GGDEF domain, synthesize c-di-GMP from two molecules of guanosine triphosphate (3). Many GGDEF domains contain a consensus GGDEF motif, which serves as the catalytic active site (A-site), and an RxxD motif, which serves as an allosteric inhibition site (I-site), located before the active site (4–6). However, not all GGDEF domain-containing proteins have a functional RxxD motif in their I-site, and the presence or absence of the RxxD motif does not definitively predict whether a DGC will be active or not (7,8). The DGCs, such as EdcCE from *Erwinia amylovora*, have been reported to regulate multiple functions, including biofilm formation, swimming motility, type III secretion, and pathogenicity (9).

PDEs, containing either an EAL or HD-GYP domain, degrade c-di-GMP into either 5’-phosphoguanylyl-(3’,5’)-guanosine (pGpG), or GMP (10,11). The catalytic activity of EAL domain proteins requires divalent cations, Mg^2+^ or Mn^2+^, and a flexible loop barrel known as loop 6 (12–14). EAL domain-containing PDEs, such as RocR and FimX from *P. aeruginosa*, catalyze a reaction that converts c-di-GMP into a linear molecule, pGpG (10,12,14,15). Other PDEs, such as those containing the HD-GYP domain, like *pggH* from *Vibrio cholerae* and Orn from *P. aeruginosa*, degrade c-di-GMP into an intermediate product, pGpG, and then hydrolyze two GMP molecules as the final product (16,17).

Bacteria possess several proteins containing GGDEF and EAL domains to adapt to complex environmental conditions. These proteins often contain additional catalytic and signal receiver or transmission domains, suggesting that they respond to environmental stimuli to control c-di-GMP levels (18,19). The concentration of c-di-GMP has been reported to be tightly spatially and temporally regulated within the cell (20,21). In *C. crescentus*, the polar localization of the DGC PopA to the old cell is controlled by c-di-GMP binding, and PopA attracts CtrA–an essential regulator for cell cycle progression–to the old cell pole (22). The DGC DcpA from *Agrobacterium tumefaciens* is required for elevated unipolar polysaccharide production and surface attachment (23). Therefore, DGCs play an important role in bacterial adaptation to various ecological niches.

*Pantoea ananatis* is a plant pathogen that causes rice grain rot, sheath rot, and onion center rot. This pathogen has gradually expanded during the growing season when the weather is hot and humid in Korea (24,25). *P. ananatis* is considered an emerging pathogen because it is known to cause disease in various important crops and infect insects and humans (26,27). In *P. ananatis* PA13, the pathogenicity is regulated by quorum sensing (QS), which is bacterial cell-to-cell communication. Exopolysaccharide (EPS) production, the hypersensitive response in tobacco, and virulence in rice are all controlled by QS (28).

Recently, it has been reported that *Pantoea* sp. YR343, isolated from the rhizosphere of *Populus deltoides*, forms biofilms along the surface of plant roots and possesses plant growth-promoting characteristics. Overexpression of DGCs in this bacterium resulted in reduced motility, increased EPS production, and enhanced biofilm formation. These DGCs were expressed during plant association, and c-di-GMP signaling may be involved in root colonization by *Pantoea* sp. YR343 (29). In *Erwinia amylovora*, the causative fire blight disease, which belongs to the same Erwiniaceae family as *Pantoea*, functional studies on DGCs and PDEs related to pathogenicity have been reported. However, functional studies on the c-di-GMP system of *P. ananatis* have not been conducted previously.

In this study, we identified proteins containing GGDEF and EAL domains using complete genome data of *P. ananatis* PA13. Moreover, we constructed mutants and overexpression strains of the genes encoding these proteins and systematically investigated their phenotypic functions in mucoid, rugose and Congo Red binding phenotypes, swimming motility, and pellicle formation. This study provides preliminary characteristics of the GGDEF and EAL domain proteins in *P. ananatis* PA13 and may help to elucidate the role of c-di-GMP signaling in this rice pathogen.

## RESULTS

### GGDEF and EAL domain-containing proteins in *P. ananatis* PA13

By analyzing the complete genome sequence of *P. ananatis* PA13 using NCBI’s conserved domain database, we identified the predicted proteins, including GGDEF and EAL domains. As a result, 26 putative c-di-GMP metabolism-related proteins were identified; their predicted conserved domains and transmembrane (TM) segments are shown in Figure 1. Of the 26 proteins, 9 proteins encode GGDEF-only domain, 5 proteins encode dual GGDEF/EAL domain, and 12 proteins encode EAL-only domain. However, the gene encoding HD-GYP domain protein was not identified in *P. ananatis* PA13. Previous studies reported highly conserved residues important for the catalytic activity, enzymatic activity, and structure of GGDEF and EAL domains. Although not all proteins are like this, the presence of these residues can predict whether the c-di-GMP metabolism-related proteins are enzymatically active (4,14).

**FIG 1.**
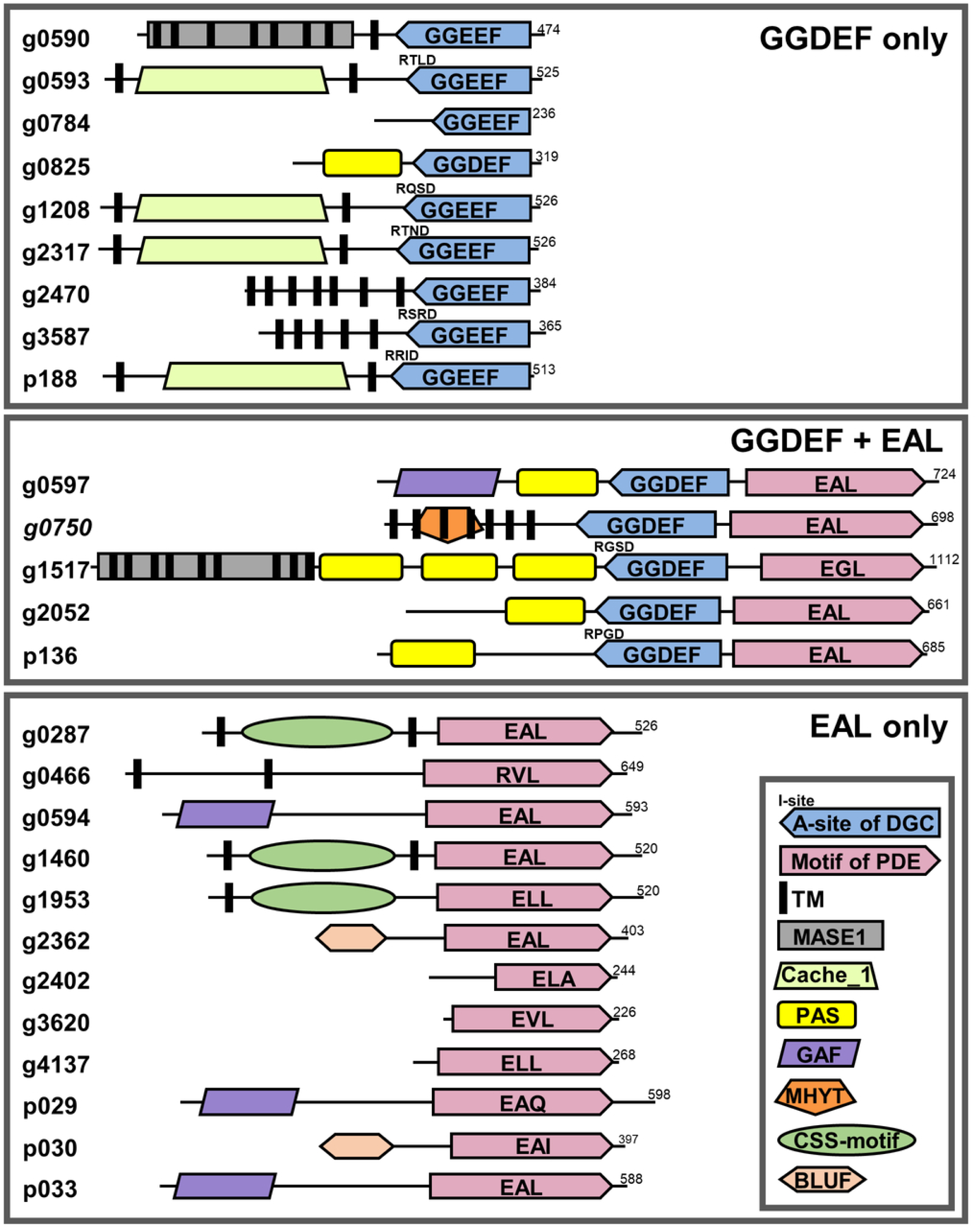
Predicted GGDEF and EAL domain architecture and conserved motifs in *Pantoea ananatis* PA13, including the possible transmembrane helices. The number indicates the size (amino acids) of the protein. GGDEF, diguanylate cyclase; EAL, phosphodiesterase; TM, transmembrane domain; MASE1, membrane-associated sensor type 1; Cache_1, small-molecules recognition domain; PAS, signal transduction domain; GAF, cGMP-specific phosphodiesterase/adenylate cyclase/FhlA; MHYT, signal transduction domain; CSS-motif, redox regulation domain; BLUF, blue light receptor domain.

Five GGDEF-only domain proteins (g0593, g1208, g2317, g3587, and p188) and 2 dual GGDEF/EAL domain proteins (g1517 and p136) were predicted to contain the consensus I-site motif, RxxD. The remaining GGDEF proteins lack a typical I-site. They have critical conserved residues for catalytic activity of GGDEF domain proteins, so they can be predicted to have enzymatic activity.

Of the 26 proteins identified, 19 proteins contain possible sensor and signal transduction domains (Fig. 1). These proteins included Per-ARNT-Sim (PAS), cGMP-specific phosphodiesterase, adenylyl cyclases, and FhlA (GAF), and conserved motif, methionine-histidine-tyrosine-threonine (MHYT), which have been regarded as signal sensors for oxygen, the cellular redox state, and potentially, other small ligands. Five proteins (g0825, g0597, g1517, g2052, and p136) contain PAS domains, regarded as signal sensors for light, redox state, and oxygen (30). Four proteins (g0597, g0594, p029, and p033) contain GAF domains, involved in protein-associated small ligand binding and protein–protein interactions (31). Four proteins (g0593, g1208, g2317, and p188) contain calcium channels and chemotaxis receptors (Cache_1) domains, involved in small-molecules recognition (32). Three proteins (g0287, g1460, and g1953) contain cysteine-rich secreted and secretion (CSS)-motif domains that allow redox regulation through disulfide bond formation (33). Two proteins (g0590 and g1517) contain membrane-associated sensor 1 (MASE1) domains, which act as sensors for the modulation of various output domain activity (34). Two proteins (g2362 and p030) contain blue light sensor using FAD (BLUF) domains, which serve as a photoreceptor in blue light-mediated signal transduction (35). The g0750 protein contains MHYT domains associated with signal transduction (36).

### Effect of overexpression of GGDEF and EAL domain protein-encoding genes on Congo Red binding, mucoid, and rugose phenotypes

To investigate the phenotypic functions of the 26 GGDEF and EAL domain-containing proteins in PA13, we constructed overexpression strains of the GGDEF and EAL domain protein-encoding genes using the isopropyl-β-D-thiogalactopyranoside (IPTG)-inducible vector pSRKGm. Each overexpression strain was assessed for Congo Red binding, mucoid and rugose phenotypes (Fig. 2). Eight GGDEF domain protein-overexpression strains (g0593, g1208, g2470, g3587, p188, g0597, g1517, and p136) exhibited colonies with a reddish color when stained with Congo red and a rougher surface than the wild type on Luria–Bertani (LB) medium, due to polysaccharide and amyloid appendages called curli (37). Five GGDEF domain protein-overexpression strains (g0825, g1208, g2470, g3587, and g1517) showed significantly reduced mucinous phenotype on Agrobacterium minimal (AB) medium supplemented with 0.2% glucose (ABG) and exhibited wrinkled colonies and 2 EAL domain protein-overexpression strains (g0594 and p029) elevated mucoid phenotypes. Five GGDEF domain protein-overexpression strains (g0593, g1208, g2470, g3587, and p136) exhibited a rugose appearance on LB medium. In particular, 3 GGDEF domain protein-overexpression strains (g1208, g2470, and g3587) showed significantly different characteristics compared to the wild type in all phenotypes, including Congo Red binding, mucoid, and rugose phenotypes. This result indicates that higher concentrations of intracellular c-di-GMP via overexpression of GGDEF domain protein-encoding genes increased the production of extracellular matrix components in PA13.

**FIG 2.**
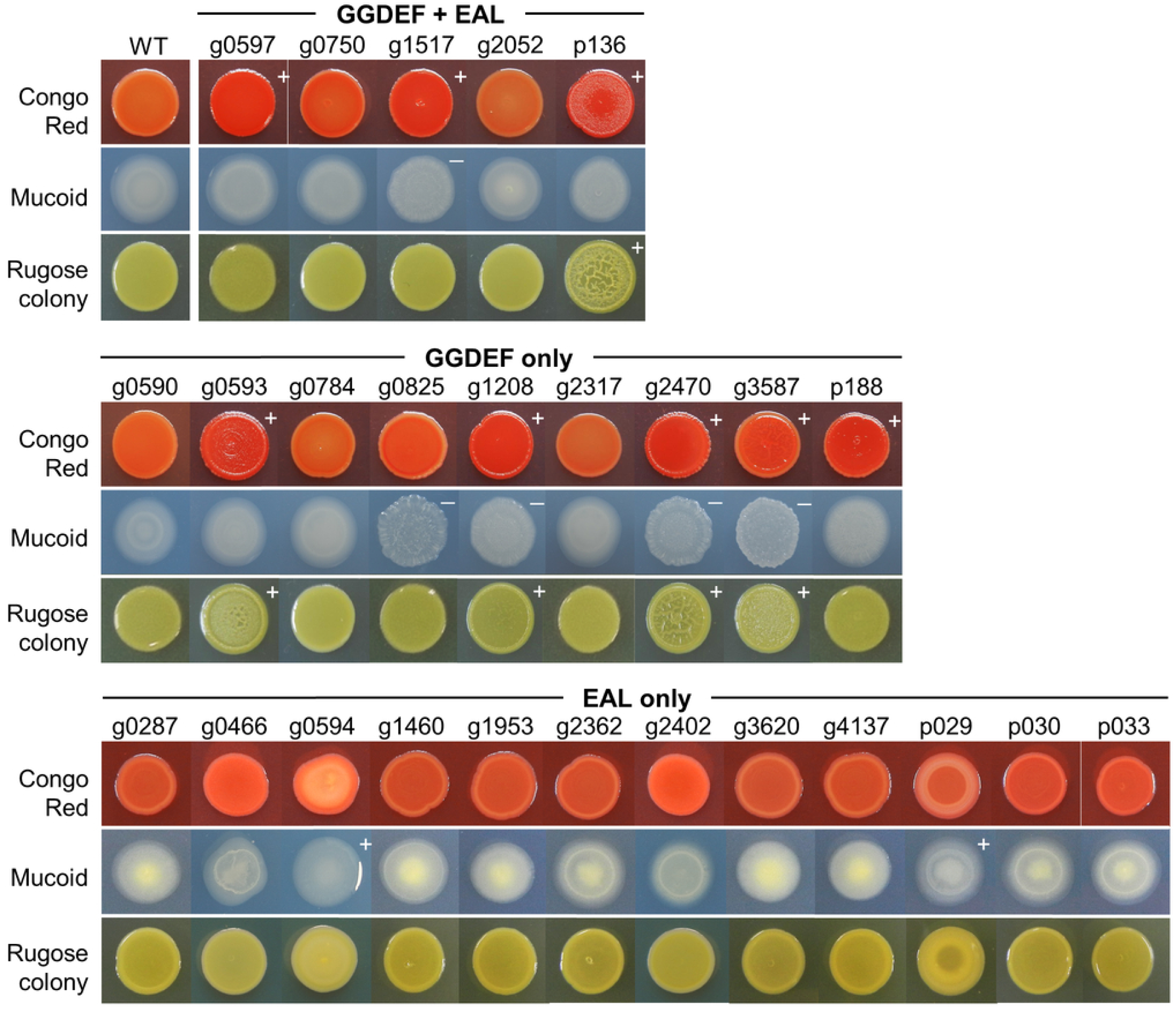
Congo Red binding, mucoid, and rugose phenotypes after overexpression of individual genes encoding GGDEF or EAL domain proteins. Congo Red binding and rugose colony morphology after 48 h, or mucoid phenotype after 24 h for the indicated wild type and gene-overexpressing strains was evaluated with 500 µM IPTG on Luria–Bertani (LB) or Agrobacterium minimal medium with 0.2% glucose (ABG), respectively. + denotes enhanced activity or rugose colony morphology; − denotes reduced activity compared to those of the wild type (WT).

### Effect of overexpression of GGDEF and EAL domain protein-encoding genes on pellicle formation

Overexpression of the gene encoding GGDEF domain protein activity induces cellulase-sensitive pellicle formation in *Burkholderia glumae* (38). We assessed the effects of GGDEF and EAL domain protein-overexpression strains on pellicle formation. Eight GGDEF domain protein-overexpression strains (g0593, g1208, g2470, g3587, p188, g0597, g1517, and p136) exhibited pellicle formation in LB medium (Fig. 3A). Interestingly, the 5 overexpression strains with pellicle formation were consistent with those that displayed red colonies in the Congo Red binding assay (Fig. 2 and 3A). This result indicates that the overexpression of GGDEF domain protein-encoding genes in PA13 affects stimulation of the extracellular matrix component, including pellicle formation. None of the EAL domain protein-overexpressing strains showed significant changes in pellicle formation compared to the wild type.

**FIG 3.**
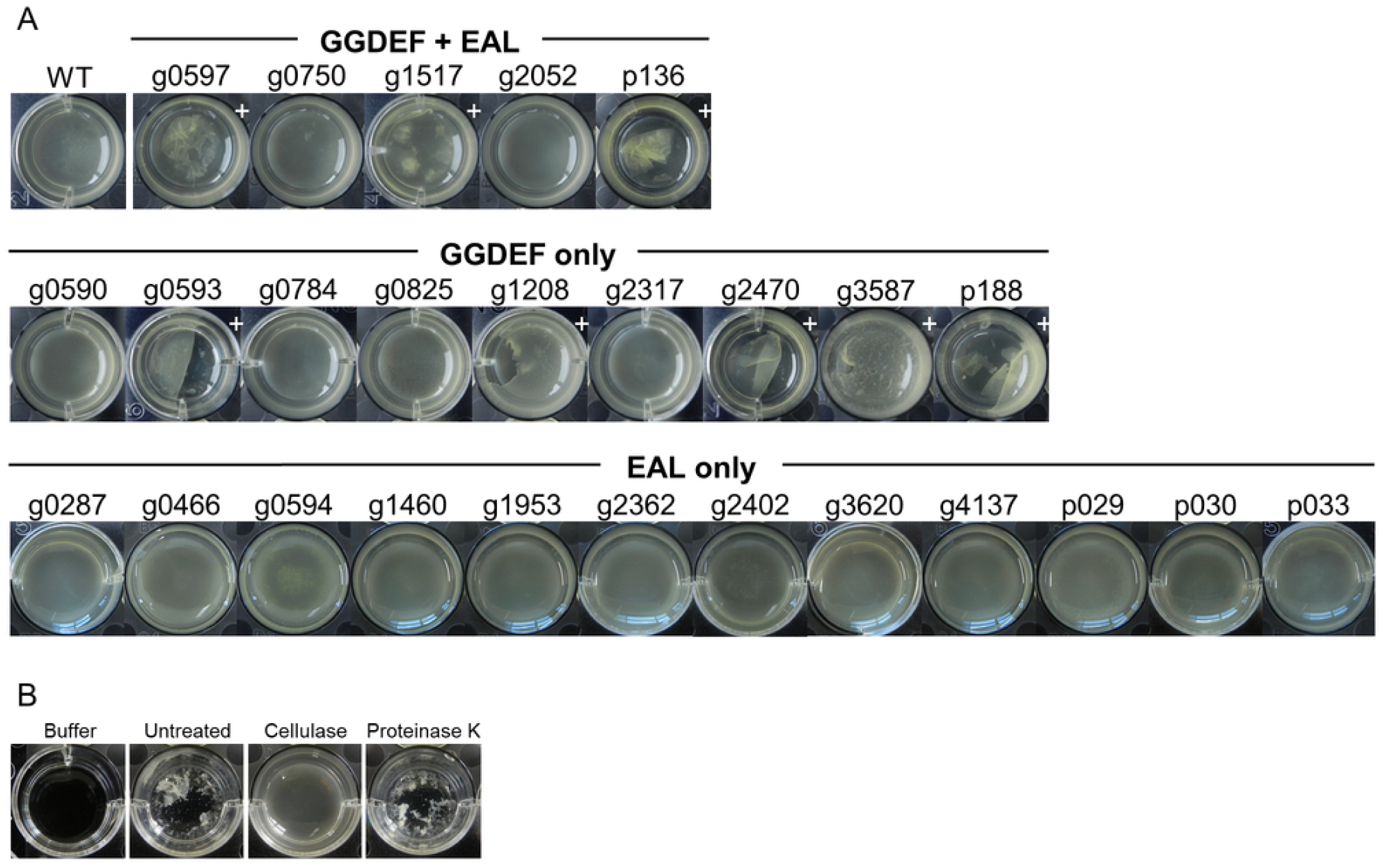
Pellicle formation after overexpression of individual genes encoding GGDEF or EAL domain proteins and pellicle degradation by cellulase. (A) Static pellicle formation for the indicated wild type (WT) and gene-overexpressing strains was observed in LB broth with 500 µM IPTG for 48 h at room temperature. + denotes pellicle formation. (B) Pellicle degradation assays. The pellicle harvested from the static culture of the g3587-overexpressing strain at room temperature was treated with cellulase or Proteinase K in phosphate-buffered saline. The pellicle was degraded by the addition of cellulase, but Proteinase K did not degrade the pellicle.

Cellulase is an enzyme that breaks down cellulose, a complex carbohydrate that forms the structural basis of bacterial pellicle (38). The enzyme hydrolyzes the β-1,4-glycosidic linkages in cellulose, leading to the breakdown of the bacterial pellicle structure (39). To determine the major constituents of the pellicle produced by *P. ananatis*, the sensitivity of the pellicle to cellulase was evaluated. When the 2-day-old pellicles produced in all strains overexpressing the GGDEF domain protein were treated with cellulase, the aggregated pellicles decomposed and became suspended, which demonstrates sensitivity to cellulase and suggests that the major components of the pellicle are substances sensitive to cellulase. Representatively, the cellulase-sensitive pellicle of the g3587-overexpressing strain and pellicle degradation by cellulase are shown in Figure 3B. No pellicle degradation by Proteinase K occurred, which was consistent with previously reported results (38).

### Effect of overexpression of GGDEF and EAL domain protein-encoding genes on swimming motility

We also assessed the effects of GGDEF and EAL domain protein-overexpression strains on swimming motility (Fig. 4). Compared to the wild type, 3 GGDEF domain protein-overexpression strains (g0825, g3587, and p188) exhibited attenuated motility by approximately 40%, 30%, and 25%, respectively. Also, 4 EAL domain protein-overexpression strains (g0466, g0594, g2402, and p029) showed attenuated motility by approximately 17%, 23%, 35%, and 26%, respectively. These results suggest that 3 EAL domain protein-overexpression strains and 4 EAL domain protein-overexpression strains negatively regulate swimming motility in PA13.

**FIG 4.**
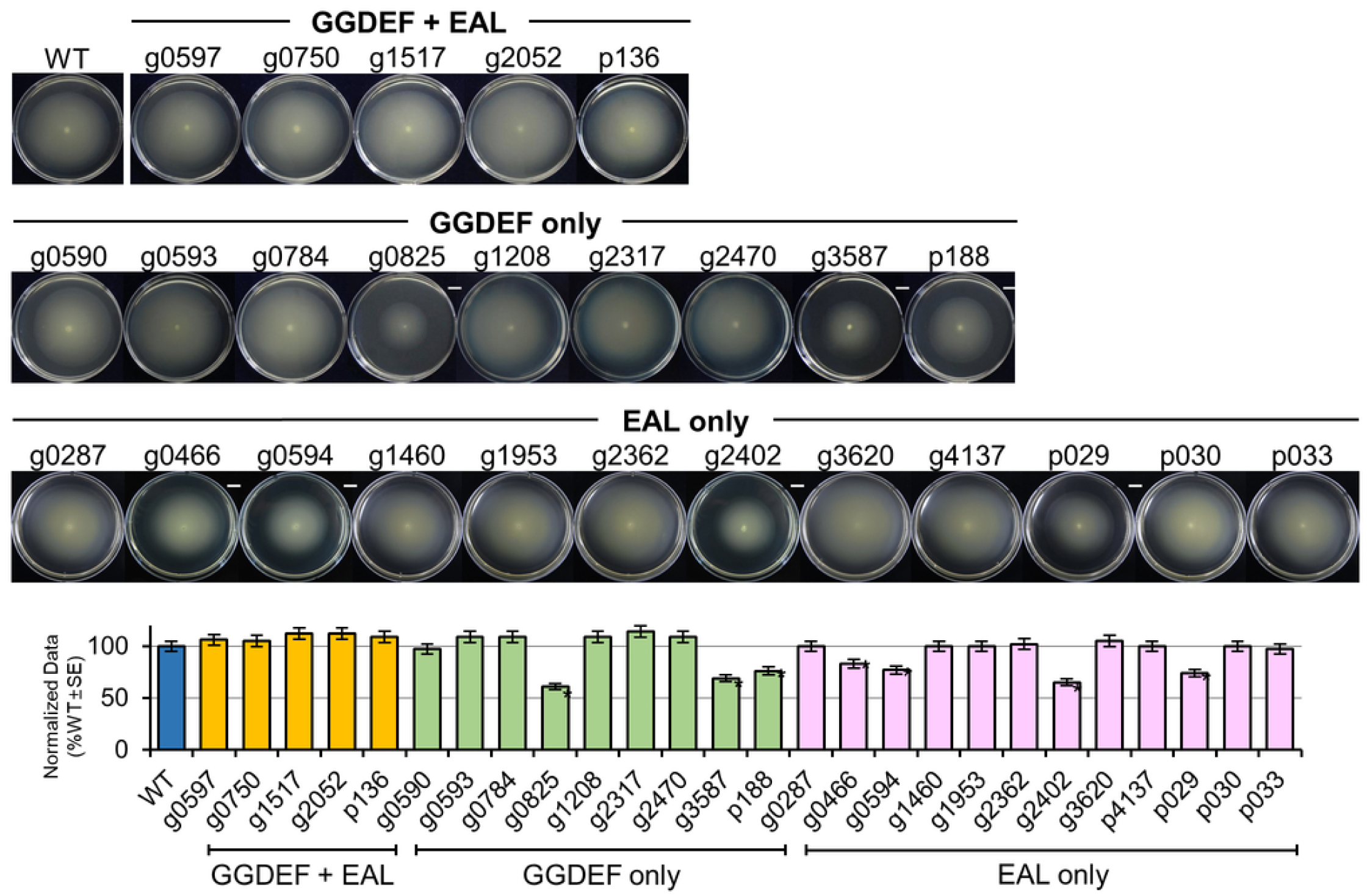
Swimming motility after overexpression of individual genes encoding GGDEF or EAL domain proteins. Swimming motility for the indicated wild type (WT) and gene-overexpressing strains was evaluated on LB plates with 0.3% Bacto agar and 500 µM IPTG after incubating at 28°C for 24 h. The swimming ring of the indicated gene-overexpressing strains was normalized to that of the WT strain. Bars are presented as means ± standard error (*n* = 3). Columns marked with an asterisk (*) indicate significant differences between the mutant and the WT strain using Student’s *t*-test (*P* ≤ 0.05). − denotes reduced swimming motility compared to those of the WT.

### Effect of mutations in the GGDEF and EAL domain protein-encoding genes on c-di-GMP-dependent phenotypes for sessile lifestyles

To determine whether mutation of 26 GGDEF and EAL domain-containing proteins affected the phenotypic functions of PA13, we constructed insertion mutants of GGDEF and EAL domain protein-encoding genes following homologous recombination via a Campbell-like mechanism (40). Each mutant was assessed for Congo Red binding, mucoid and rugose phenotypes, pellicle formation, and swimming motility (Table 1). Only the EAL domain protein-encoding gene, g2052::pCOK238 mutant exhibited colonies with a reddish color on LB medium supplemented with Congo Red and pellicle formation in LB medium (Fig. 5). No other mutants had significant changes in Congo Red binding or pellicle formation compared to the wild type. We therefore designated the g2052 gene as *pdeC* that regulates Congo Red binding and cellulase-sensitive pellicle formation phenotypes to distinguish it from other putative EAL domain motif genes. For complementation, plasmid pYS46 was constructed by cloning the full-length g2052 (*pdeC*) gene into pSRKGm, and the mutant transformed with pYS46 restored Congo Red binding and pellicle formation to wild type levels in the presence of 500 µM IPTG (Fig. 5). Overexpression of the g2052 (*pdeC*) gene with both GGDEF and EAL domains exhibited no phenotype alterations, including Congo Red binding or mucoid phenotype (Fig. 2), whereas the mutant induced enhanced Congo Red binding and pellicle formation (Fig. 5). These results suggest that g2052 (PdeC) has EAL enzymatic activity rather than GGDEF enzymatic activity.

**FIG 5.**
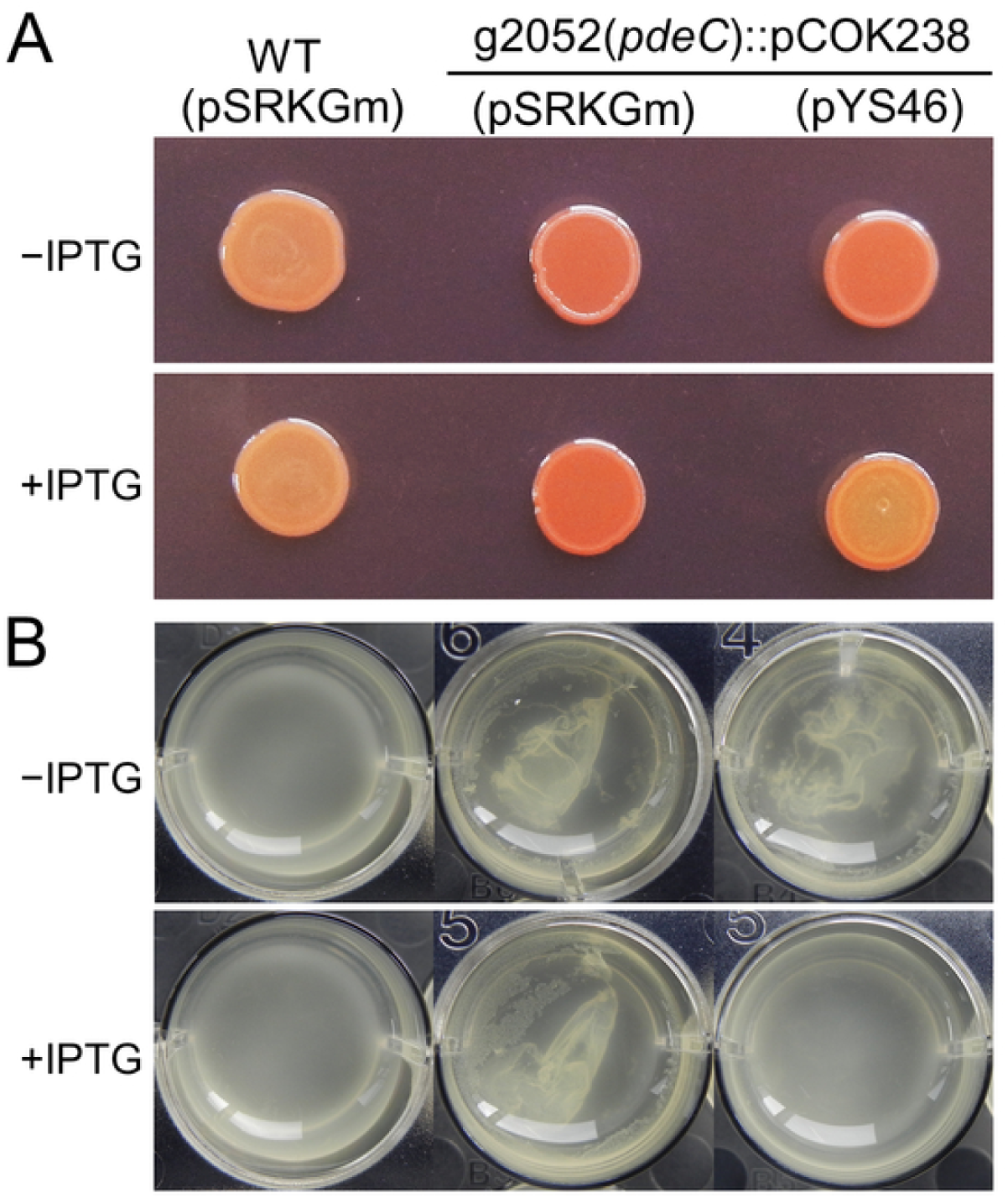
Congo Red binding phenotype and pellicle formation of g2052 mutants encoding EAL domain proteins. Congo Red binding (A) and pellicle formation (B) after 48 h for the indicated wild type (WT) and mutant strains with (+) or without (−) 500 µM IPTG.

**TABLE 1.** Congo Red binding, pellicle formation, mucoid and rugose phenotypes, swimming motility, and virulence of overexpression strains and mutants of 26 GGDEF and EAL domain-containing proteins.

| Strain | DGC or PDE motif | Phenotypes |  |  |  |  |  |
| --- | --- | --- | --- | --- | --- | --- | --- |
|  |  | Congo Red binding | Pellicle formation | Mucoid colony | Rugose colony | Swimming motility | Virulence |
| Wild type |  | + | – | + | – | + | + |
| Overexpression strains |  |  |  |  |  |  | + |
| g0590 | GGDEF | + | – | + | – | + | + |
| g0593 | GGDEF | ++ | + | + | + | + | + |
| g0784 | GGDEF | + | – | + | – | + | + |
| g0825 | GGDEF | + | – | – | – | – | + |
| g1208 | GGDEF | ++ | + | – | + | + | + |
| g2317 | GGDEF | + | – | + | – | + | + |
| g2470 | GGDEF | ++ | + | – | + | + | + |
| g3587(*dgcM*) | GGDEF | ++ | + | – | + | – | + |
| p188 | GGDEF | ++ | + | + | – | – | + |
| g0597 | GGDEF+EAL | ++ | + | + | – | + | + |
| g0750 | GGDEF+EAL | + | – | + | – | + | + |
| g1517 | GGDEF+EGL | ++ | + | – | – | + | + |
| g2052 | GGDEF+EAL | + | – | + | – | + | + |
| p136 | GGDEF+EAL | ++ | + | + | + | + | + |
| g0287 | EAL | + | – | + | – | + | + |
| g0466 | RVL | + | – | + | – | – | + |
| g0594 | EAL | + | – | ++ | – | – | + |
| g1460 | EAL | + | – | + | – | + | + |
| g1953 | ELL | + | – | + | – | + | + |
| g2362 | EAL | + | – | + | – | + | + |
| g2402 | ELA | + | – | + | – | – | + |
| g3620 | EVL | + | – | + | – | + | + |
| g4137 | ELL | + | – | + | – | + | + |
| p029 | EAQ | + | – | ++ | – | – | + |
| p030 | EAI | + | – | + | – | + | + |
| p033 | EAL | + | – | + | – | + | + |
| Mutants |  | + | – | + | – | + | + |
| g0590::pCOK532 | GGDEF | + | – | + | – | + | + |
| g0593::pCOK533 | GGDEF | + | – | + | – | + | + |
| g0784::pCOK534 | GGDEF | + | – | + | – | + | + |
| g0825::pCOK535 | GGDEF | + | – | + | – | + | + |
| g1208::pCOK536 | GGDEF | + | – | + | – | + | + |
| g2317::pCOK537 | GGDEF | + | – | + | – | + | + |
| g2470::pCOK538 | GGDEF | + | – | + | – | + | + |
| g3587::pCOK516 | GGDEF | + | – | + | – | + | + |
| p188::pCOK539 | GGDEF | + | – | + | – | + | + |
| g0597::pCOK246 | GGDEF+EAL | + | – | + | – | + | + |
| g0750::pCOK247 | GGDEF+EAL | + | – | + | – | + | + |
| g1517::pCOK248 | GGDEF+EGL | + | – | + | – | + | + |
| g2052(*pdeC*)::pCOK238 | GGDEF+EAL | ++ | + | + | – | + | + |
| p136::pCOK249 | GGDEF+EAL | + | – | + | – | + | + |
| g0287::pCOK243 | EAL | + | − | + | − | + | + |
| g0466::pCOK244 | RVL | + | − | + | − | + | + |
| g0594(*pdeM*)::pCOK239 | EAL | + | − | − | − | + | + |
| g1460::pCOK250 | EAL | + | − | + | − | + | + |
| g1953::pCOK240 | ELL | + | − | + | − | + | + |
| g2362::pCOK241 | EAL | + | − | + | − | + | + |
| g2402(*pdeS*)::pCOK251 | ELA | + | − | + | − | ++ | + |
| g3620::pCOK254 | EVL | + | − | + | − | + | + |
| g4137::pCOK252 | ELL | + | − | + | − | + | + |
| p029::pCOK245 | EAQ | + | − | + | − | + | + |
| p030::pCOK253 | EAI | + | − | + | − | + | + |
| p033::pCOK255 | EAL | + | − | + | − | + | + |

A decreased or increased phenotype relative to wild type is indicated by one of −, +, or ++.

Only the EAL domain protein-encoding gene, g0594::pCOK239 mutant exhibited a non-mucoid phenotype on ABG medium compared to the wild type, and no other traits were changed. No other mutants had a significant change in mucoid phenotype. Therefore, we designated the g0594 gene to *pdeM*, which regulates mucoid phenotype, to distinguish it from other putative EAL domain motif genes. A complementation strain transformed with pCOK286/pSRKGm::g0594(*pdeM*) exhibited a hyper-mucinous phenotype in the presence of 500 µM IPTG (Fig. 6). This is consistent with the overexpression of the g0594 gene with the EAL domain elevated mucoid phenotype (Fig. 2). These results indicate that the EAL-only domain protein g0594 (PdeM) positively regulates the mucoid phenotype of *P. ananatis* PA13.

**FIG 6.**
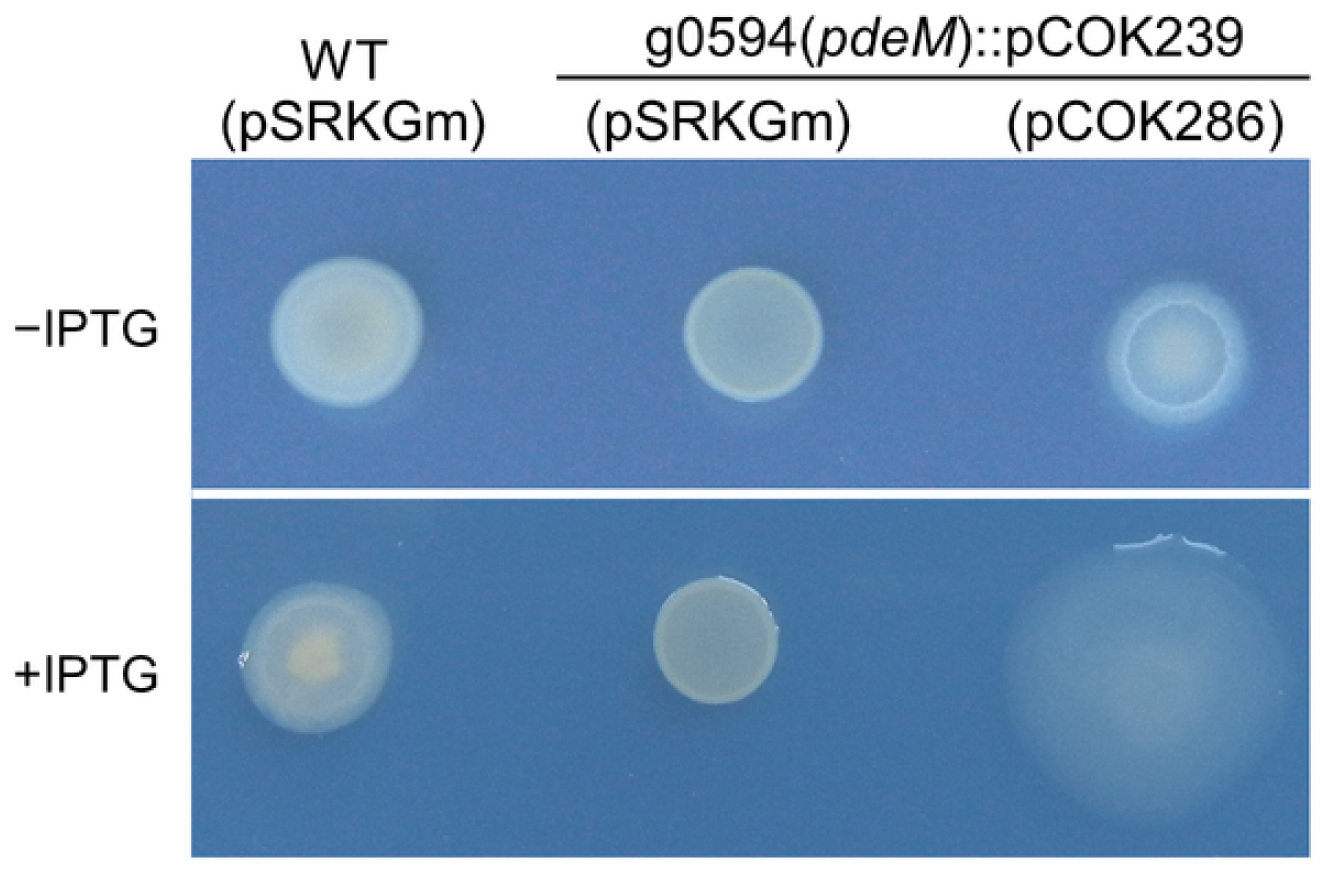
Mucoid phenotype of g0594 mutants encoding EAL domain proteins. Mucoid phenotype after 24 h for the indicated wild type (WT) and mutant strains on ABG agar plates with (+) or without (−) 500 µM IPTG.

Only the EAL domain protein-encoding gene, g2402::pCOK251 mutant significantly enhanced swimming motility by 29% compared to the wild type. No other mutants showed significant changes in swimming motility compared to the wild type. Therefore, we designated the g2402 gene to *pdeS*, which regulates swimming motility, to distinguish it from other putative EAL domain motif genes. A complementation strain transformed with pCOK287/pSRKGm::g2402(*pdeS*) reduced the swimming motility in the presence of 500 µM IPTG, which was significantly reduced by 28% compared to the wild type (Fig. 7). This is consistent with the overexpression of the g2402 gene with the EAL domain reduced swimming motility (Fig. 4). The three genes (g0466, g0594, and p029), whose swimming motility was attenuated by overexpression, exhibited no change by a mutation in swimming motility (Fig. 4 and Fig. 7). These results indicate that the EAL-only domain protein g2402 (PdeS) negatively regulates swimming motility in *P. ananatis* PA13.

**FIG 7.**
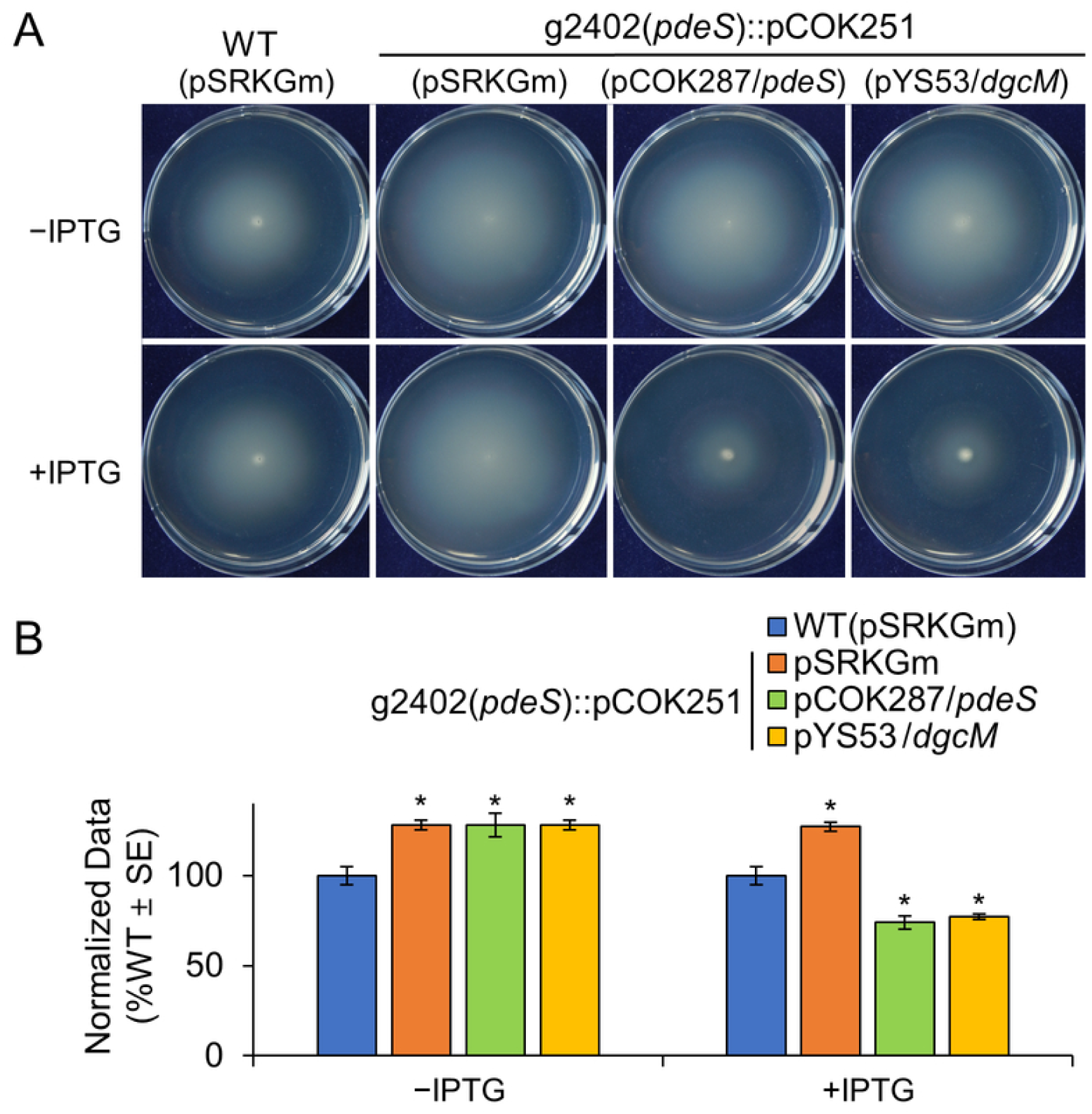
Swimming motility of g2402 mutants encoding EAL domain proteins. (A) Swimming motility after 24 h for the indicated wild type (WT) and mutant strains with (+) or without (−) 500 µM IPTG. (B) Quantification of swimming ring diameters was measured after 24 h on LB plates with 0.3% Bacto agar at 28°C. Data for swimming ring diameters were the mean of three independent experiments (*n* = 3). For presentation, all data were normalized to results obtained with the WT and are expressed as %wild type ± standard error. Columns marked with an asterisk (*) indicate significant differences between the mutant and the WT strain using Student’s *t*-test (*P* ≤ 0.05).

In *E. coli*, YdiV (237 amino acids), an EAL-only domain protein, inhibits motility through its interaction with FlhD to abolish FlhDC interaction with DNA (41,42). Since PdeS (244 amino acids) of PA13 is also an EAL-only domain protein and its mutation leads to increased motility, we hypothesize that PdeS functions similarly to YdiV. To analyze this, a phylogenetic analysis of EAL domains from both multidomain and EAL-only domains revealed that PdeS clusters closely with YdiV, distinct from multidomain EAL proteins (Fig. 8A). Furthermore, we compared the domain similarity of PdeS from PA13 with YdiV homologs from *Salmonella enterica serovar* Typhimurium ATCC 14028, *E. coli* K-12, *Shigella flexneri* 2a, *Klebsiella pneumoniae*, and *Yersinia enterocolitica* (41,43). Consistent with YdiV in other enterobacteria, PdeS possesses conserved active sites for PDE activity, a dimerization loop, and FlhD-binding residues (Fig. 8B). Collectively, these results suggest that PdeS of PA13 is an EAL-only protein that has the same functional role as YdiV.

**FIG 8.**
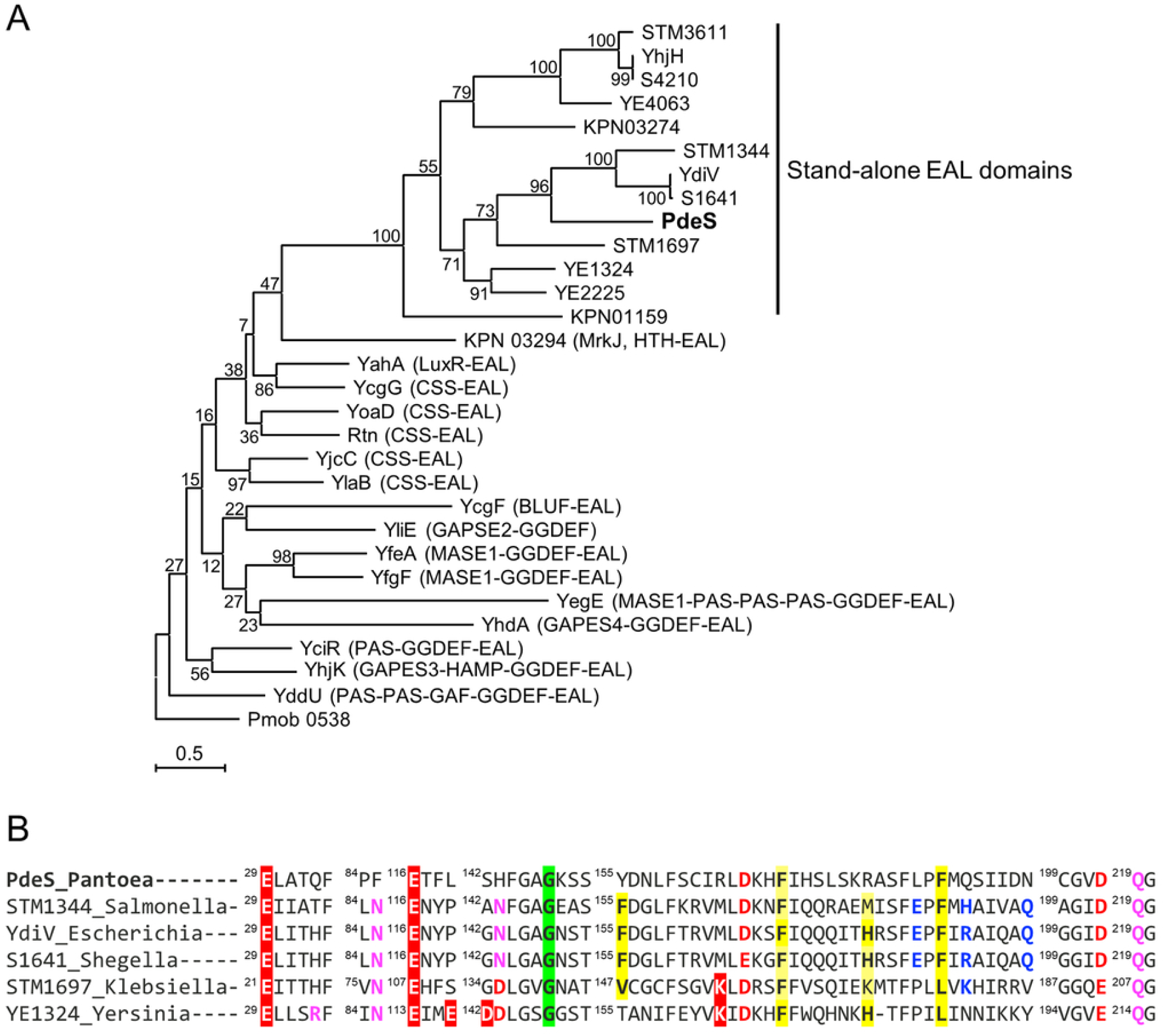
Phylogenetic tree of EAL domain proteins and conserved amino acid residues of EAL-only proteins. (A) The phylogenetic tree was generated under the most suitable substitution model, evaluated automatically, while branch support was estimated using the Bayes algorithm. Phylogenetic relationship of enterobacterial EAL domains and PdeS, inferred by maximum likelihood, according to the report previously described (41,43). The PdeS protein of *P. ananatis* PA13 is in bold. (B) Amino acid sequence alignment for the conserved active site, dimerization loop, and FlhD-binding residues of the PdeS among EAL-only proteins as previously described (41,43). Proteins from *Salmonella enterica* serovar Typhimurium ATCC 14028, *E. coli* K-12, *Shigella flexneri* 2a, *Klebsiella pneumoniae*, and *Yersinia enterocolitica* are listed by their respective names in the NCBI protein database. The residues forming the active site of c-di-GMP PDE are shown in white on a red background. Residues involved in the dimerization loop are shaded in green. Conserved residues located near the active site are colored red or magenta. Hydrophobic residues involved in FlhD binding are shaded in yellow, and charged residues at the interface are shown in blue.

Next, we transferred the multi-copy number plasmid pYS53 carrying an intact g3587 gene to the g2402(*pdeS*)::pCOK251 mutant to determine whether the g3587 gene plays a key role in c-di-GMP-dependent sessile phenotypes. The enhanced swimming motility of the g2402(*pdeS*)::pCOK251 mutant was reduced by transformation of pYS53, and the decrease was comparable to that of the mutant transformed with pCOK287 (Fig. 7). These results indicated that a putative GGDEF domain protein gene (g3587) possessing TM segments was the most influential for c-di-GMP-dependent sessile phenotypes, including motility, when constitutively expressed in the multi-copy number plasmid pYS53. Therefore, we designated the g3587 gene as the master GGDEF domain protein, *dgcM*, to distinguish it from other putative GGDEF domain protein motif genes.

### Quantification of cellular c-di-GMP

To determine the levels of cellular c-di-GMP in strains with altered phenotypes due to the overexpression and mutation of GGDEF and EAL proteins, we quantified the concentration of c-di-GMP from these strains using high-performance liquid chromatography. The c-di-GMP concentrations of the strains are shown in Table 2. The c-di-GMP concentrations of all 13 GGDEF overexpression strains and 3 EAL mutants were significantly increased compared to the wild type (Table 2). Notably, the overexpression of gene g0825 increased c-di-GMP concentrations to the millimolar range, consistent with our results indicating that g0825 functions as a GGDEF domain protein. The overexpression of the g3587 protein, which has GGDEF domain protein function and affects multiple phenotypes, also resulted in a significantly higher concentration of c-di-GMP compared to the wild type. These results suggest that these proteins play an enzymatic role in the production of c-di-GMP.

**TABLE 2.**
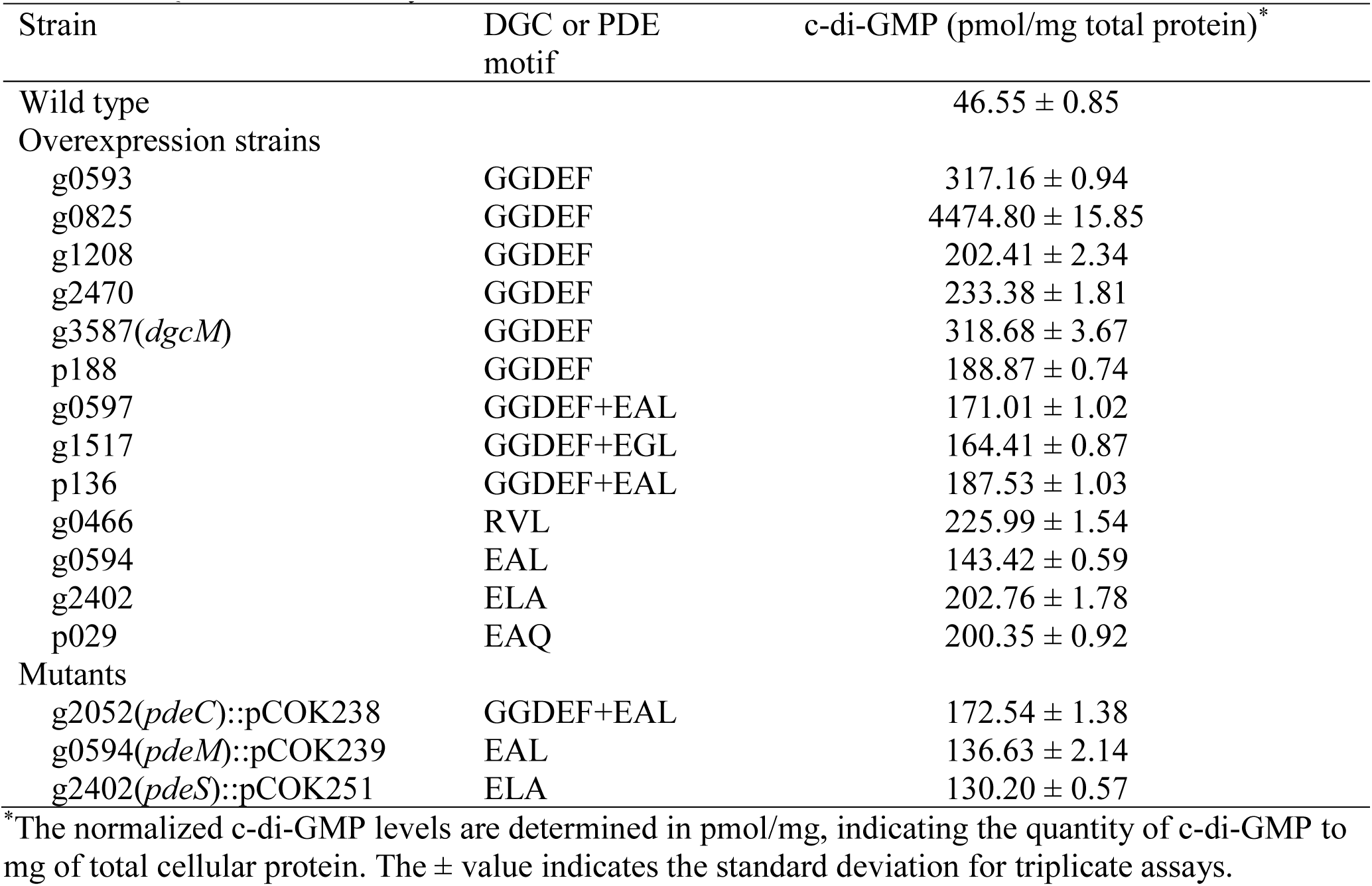
Quantification of cyclic-di-GMP in *P. ananatis*.

### Enzymatic activity prediction based on the c-di-GMP concentrations and phenotypic observation of DGC- and PDE-encoding genes

Based on the cellular c-di-GMP concentrations and the phenotypic observation of overexpression and mutant strains of c-di-GMP metabolism-related genes, 14 out of 26 proteins had a function; their roles are summarized in Figure 9. Specifically, 6 GGDEF-only domain proteins (g0593, g0825, g1208, g2470, g3587 [DgcM], and p188; blue), 4 dual GGDEF/EAL domain proteins (g0597, g1517, g2052, and p136), and 4 EAL-only domain proteins (g0466, g0594, g2402 [PdeS] and p029 ; pink) altered phenotypes, such as Congo Red binding, mucoid and rugose phenotypes, pellicle formation, and swimming motility. Through phenotypic observation, g0597, g1517, and p136 of the dual GGDEF/EAL domain proteins exhibited GGDEF enzymatic activity (blue), and g2052 (PdeC) exhibited EAL enzymatic activity (pink). EAL domain protein g0594 (PdeM) was involved only in the mucoid phenotype and not in other c-di-GMP-dependent phenotypes (Fig. 9A). All overexpression strains and mutants of GGDEF/ EAL domain proteins that exhibited a phenotypic change compared to the wild type had a higher cellular c-di-GMP level than the wild type based on HPLC detection (Table 2).

**FIG 9.**
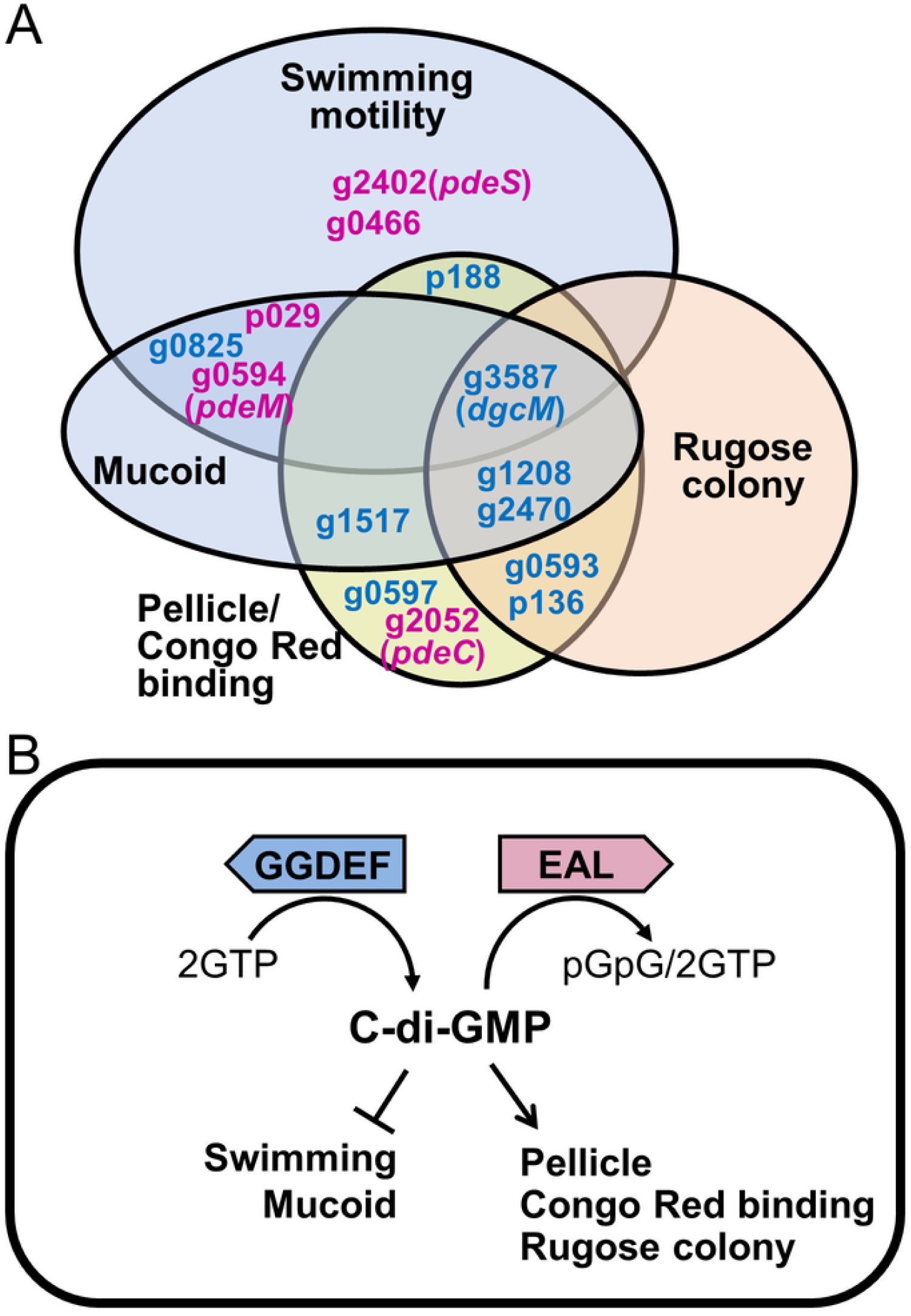
Summary of GGDEF/EAL domain-containing proteins in *P. ananatis* PA13. (A) Overlapping and discrete regulatory actions of GGDEF/EAL domain proteins on mucoid and rugose colony, cellulase-sensitive pellicle formation/Congo Red binding, and swimming motility. (B) Function of c-di-GMP in *P. ananatis* PA13. The turnover of c-di-GMP, controlled by diguanylate cyclase GGDEF proteins and phosphodiesterase EAL, positively or negatively regulates Congo Red binding, mucoid and rugose phenotypes, pellicle formation, and motility. The proteins exhibiting GGDEF enzymatic activity are marked in blue. The protein exhibiting EAL enzymatic activity are marked in pink. Arrows indicate activation of the phenotypes whereas blocking lines indicate inhibition of the phenotype.

Many GGDEF domain proteins had overlapping effects on pellicle/cellulose production, Congo Red binding, mucoid and rugose phenotypes, and swimming motility, but only g3587 (*dgcM*) affected all four phenotypes (Fig. 9A). Our results show that increased c-di-GMP concentrations decreased swimming motility and mucoid, increased pellicle formation, and changed colony morphology to rugose in *P. ananatis* (Fig. 9B). In many other bacteria, c-di-GMP signaling has also been shown to inhibit motility, change colony morphology, and increase biofilm formation (44–48).

## DISCUSSION

It is well known that c-di-GMP serves as a crucial second-messenger molecule in various physiological processes in bacteria, particularly those related to the motile-to-sessile lifestyle switch, in bacteria, including plant pathogens (3,9,11,22,45). This study aimed to identify proteins that regulate c-di-GMP levels in *P. ananatis* PA13, a pathogenic bacterium that infects rice and onion, and to systematically and phenotypically investigate their functions. Previous studies have shown that the expression levels of GGDEF and EAL-encoding genes are often low or not definitively revealed, and changes in the cellular c-di-GMP concentrations of mutant strains can be difficult to detect (47). Therefore, we designed overexpression and mutation experiments to identify potential phenotypes and enzymatic activities controlled by GGDEF and EAL domain-encoding genes. We systematically investigated the morphological and physiological changes of the overexpression and mutant strains of 26 GGDEF and EAL domain-encoding genes in PA13 and measured intracellular c-di-GMP concentrations.

Overall, most DGC-encoding genes in PA13 exhibited significant phenotypic changes under overexpression but not upon mutation. Among the PDE-encoding gene mutants, only some displayed distinct phenotypic changes. The intracellular c-di-GMP concentrations were significantly elevated in the overexpression strains and mutants showing phenotypic changes compared to the wild type. These results suggest that overexpressing DGC and PDE-encoding genes is a promising strategy for investigating phenotypic changes in *P. ananatis* PA13. Although gene overexpression does not represent a natural phenomenon, it appears to be an effective method for predicting bacterial responses to specific environmental conditions.

Overexpression of DGC and PDE-encoding genes in PA13 significantly altered various cellular phenotypes. Among the many DGCs in PA13, single gene overexpression often simultaneously affected 1 to as many as 5 colony phenotypes, including Congo red binding, rugose colony morphology, mucoid colony morphology, and pellicle formation (Table 1). Outer membrane and surface properties including the presence of adhesive structures such as extracellular cellulose and amyloid curli fibers are the main factors that increase Congo red binding (48). Congo red binding and rugose morphotypes are directly promoted by c-di-GMP concentration in both *Salmonella enterica* serovar *Typhimurium* and *Vibrio cholerae* (49,50). In *Acetobacter xylinum* (now *Gluconacetobacter xylinus*), it was first reported that c-di-GMP is a positive effector that activates cellulose synthase (51). In *E. coli*, the loss of the GGDEF-domain gene *ydaM* specifically reduced Congo red binding and serves as a critical regulator of CsgD, which controls curli and cellulose production (52). Similarly, c-di-GMP levels affect pellicle formation in *P. aeruginosa* (53). The Pel pathway, present in diverse microbes, synthesizes and exports Pel polysaccharides that play an important role in pellicle and biofilm formation (54).

Previous studies have demonstrated that c-di-GMP signaling proteins influence EPS production in *P. aeruginosa* PA14 (54, 55). In *Erwinia amylovora*, c-di-GMP positively regulates biofilm formation and amylovoran, an acidic polysaccharide composed of repeating units of galactose and glucuronic acid (9). In PA13, we also identified 7 DGC and PDE-encoding genes (g1517, g0825, g1208, g2470, g3587, g0594, and p029) that affect EPS production (Table 1, Fig. 2). Based on the genome analysis of PA13, an EPS capsule biosynthetic operon exists, and its genes are predicted to be directly or indirectly regulated by c-di-GMP, warranting further investigation.

Overexpression of 8 DGC and PDE-encoding genes (g0825, g3587, p188, g0426, g0594, g0594, g2402, and p029) in PA13 significantly reduced motility. While high intracellular c-di-GMP levels typically inhibit bacterial motility, the *pdeS* mutant of *P. ananatis* PA13 exhibited an enhanced motility phenotype (Fig. 7). Given that the EAL-domain protein YdiV acts as a flagellar repressor, the high structural and functional similarity between PdeS and YdiV (Fig. 8) suggests that PdeS plays a homologous FlhD inhibitory role. These findings identify PdeS as a putative flagellar gene inhibitor in PA13. In *Salmonella* and *E. coli*, YdiV has been shown to repress FlhD under nutrient-starvation or iron-limiting conditions, thereby inhibiting flagellar synthesis and reducing motility (43,56). Future studies need to focus on determining the precise conditions under which PdeS is expressed and regulated.

Domain analysis of 26 DGC and PDE proteins in PA13 revealed that 19 contain various signal response domains, suggesting they synthesize or degrade c-di-GMP in response to environmental signals. Additionally, 13 of the 26 proteins possess N-terminal transmembrane (TM) regions, indicating localization to cell membrane. Overexpression of g3587 (*dgcM*), which contain five TM segments, triggered all c-di-GMP-dependent sessile phenotype changes, including increased Congo Red binding, conversion to rugose colony morphology and pellicle formation, and reduced mucoid phenotype and swimming motility (Figs. 1–4), indicating that the g3587 (*dgcM*) gene suggests a key role in c-di-GMP-dependent sessile phenotypes. The g3587 (*dgcM*) gene lacks any N-terminal sensor or signal transduction domain and contains only a single GGDEF domain and five TM segments, suggesting that it may localize to the membrane, sense environmental changes, interact with other adapter proteins, and exhibit enzymatic activity. These results suggest that further investigation of the protein–protein interactions of DgcM is needed (Fig. 9A). Both the g0593-overexpressing strain with two TM segments and the p136-overexpressing strain with the PAS domain exhibited identical phenotypes to each other, including in Congo Red binding, rugose colony morphology, and pellicle formation (Fig. 9A). Similarly, overexpression of g1208 and g2470, which contain 2 and 7 TM segments, respectively, induced the same c-di-GMP-dependent phenotypes, including increased Congo Red binding, conversion to rugose colony morphology, and pellicle formation, and reduced mucoid phenotype (Table1, Fig. 9A). These findings indicate that multiple DGC and PDE proteins in PA13 redundantly affect the same phenotype, such as colony morphology. Furthermore, these results suggest a potential functional link between these proteins, although further studies are required for confirmation. Our observations are consistent with previous reports showing that c-di-GMP signaling proteins do not all exhibit enzymatic activity under the same conditions; rather, specific GGDEF or EAL domain proteins are activated only under specific temporal and signaling conditions (20,21,57).

It has been reported that elevated intracellular c-di-GMP levels attenuate virulence in many plant pathogens (58,61). To investigate the role of the c-di-GMP metabolism-related genes in PA13 virulence, GGDEF-overexpressing strains and EAL mutants were inoculated into onion bulbs by the infiltration method. All overexpressing strains and EAL mutants caused onion center rot at levels comparable to the wild type (data not shown). These results suggest that the c-di-GMP signaling genes in PA13 are not associated with pathogenicity; however, the situation may be more complex. One possible explanation is that, because the infiltration method bypasses early invasion steps, the inoculated bacteria may not have experienced the effects of altered c-di-GMP metabolism during the initial stage of infection. In *P. syringae* pv. *tomato* DC3000, the c-di-GMP overexpression strain exhibited reduced virulence in *Arabidopsis thaliana* under spray inoculation but remained virulent under infiltration inoculation. This result demonstrated that the pathogen directly invaded the plant through infiltration inoculation, eliminating the adverse effect of c-di-GMP overexpression and thereby causing disease (59,62). In *Erwinia amylovora*, elevated levels of c-di-GMP conferred by the overexpression of DGC or deletion of PDE positively regulate amylovoran secretion and biofilm formation, and negatively regulate flagellar swimming motility and virulence (9,63). Consequently, the specific relationship between c-di-GMP levels and pathogenicity in PA13 requires further study.

In summary, we primarily focused on identifying the DGC and PDE-encoding genes and characterizing their phenotypic roles in PA13. We identified 26 genes involved in c-di-GMP metabolism and found 14 of them affected colony morphology and swimming motility in PA13. We also found that the c-di-GMP concentrations were significantly higher in overexpression strains or mutants of these 14 genes than in the wild type. Overall, increasing c-di-GMP levels in PA13 reduced motile-associated phenotypes and enhanced sessile phenotypes. Because c-di-GMP metabolism-related proteins are regulated in response to environmental signals, it will be important to examine their expression patterns under diverse environmental conditions and to clarify the role of c-di-GMP in PA13 pathogenesis. This study provides a foundation for understanding how the second messenger c-di-GMP functions across the environments in which plant pathogens reside.

## MATERIALS AND METHODS

### Bacterial strains and growth conditions

The bacterial strains and plasmids used in this study are listed in Table S1. *Escherichia coli* strains were cultured on Luria-Bertani (LB) medium at 37°C. *P. ananatis* PA13 strains were cultivated at 28°C on LB medium or Agrobacterium minimal medium supplemented with 0.2% glucose (ABG) medium. Antibiotics were used at the following concentrations: ampicillin, 100 µg/mL; kanamycin, 50 µg/mL; rifampicin, 50 µg/mL; gentamycin, 25 µg/mL. 5-Bromo-4-chloro-3-indoyl-b-D-galactopyranoside (X-gal) was used at 40 µg/mL when necessary. IPTG was used at 500 µM when necessary.

### DNA manipulation and data analyses

DNA manipulation, cloning, restriction enzyme digestion, and agarose gel electrophoresis were performed as previously described (64). DNA sequencing was performed by Macrogen (Seoul, Republic of Korea). DNA sequences were analyzed using the BLAST program at the National Center for Biotechnology Information (65) and the MegAlign (DNAStar, Madison, WI, USA) and GENETYX-Win (Genetyx, Tokyo, Japan) software.

Sequences of EAL domains were taken from reference 41 or GenBank/EMBL database. Sequence alignments of EAL domains for a phylogenetic analysis were generated using MEGA11. A phylogenetic tree was constructed using the maximum likelihood method with the Bayes algorithm (66).

### Conserved domain organization analysis

The conservation and functional annotation of genes encoding c-di-GMP turnover enzymes in the complete genome sequences of *P. ananatis* PA13 (GenBank accession numbers CP003085 and CP003086) were analyzed using the BLAST program (67). The predicted conserved domains of proteins were searched using BLASTP. Alignment of relevant sequence was performed using online software available at the Clustal Omega website (http://ebi.ac.uk). The presence of TM helices on *P. ananatis* PA13 proteins was predicted using TMHMM2.0 (68).

### Generation of Campbell integration

Integration of pVIK112 containing a host DNA fragment into the chromosome was performed following Campbell-type recombination (41). An internal DNA fragment of individual genes was amplified using the corresponding primer pairs (Table S2). The partial DNA fragment was purified, cloned into pGEM-T Easy (Promega, Madison, WI, USA), and confirmed by sequencing. For homologous recombination, the *Eco*RI/*Kpn*I-digested DNA fragment was cloned into the pVIK112 suicide vector. The resulting plasmids were introduced into *P. ananatis* PA13 by conjugation, and mating cells were spread on an LB medium containing kanamycin and rifampicin. The mutants were confirmed through PCR using a primer that anneals upstream of the truncated fragment and the primer LacFuse (Table S2), followed by sequencing.

### Gene overexpression

To generate the overexpression strains, we amplified each intact target gene with the primer pair (Table S2) and cloned it into the IPTG-inducible vector pSRKGm (69). The resulting plasmids were transferred into *P. ananatis* PA13 by conjugation, and mating cells were spread on LB medium containing gentamycin and rifampicin.

### Gene complementation

To generate target gene complemented strains, we amplified and cloned each intact target gene into the IPTG-inducible vector pSRKGm, generating pYS46(pSRKGm::g2052/*pdeC*), pYS53(pSRKGm::g3587/*dgcM*), pCOK286(pSRKGm::g0594/*pdeM*), and pCOK287(pSRKGm::g2402/*pdeS*), which were transferred to the corresponding mutant strains by conjugation (Table S1 and S2).

### Congo Red binding, mucoid and rugose phenotypes

Congo Red binding assay was performed on LB agar plates containing 40 µg/mL Congo Red; 5-μL overnight-cultured bacterial suspensions were spotted onto the plates and then incubated at 28°C for 48 h. The color changes of colonies were investigated. The mucoid phenotype was determined as previously described (70); 5-μL overnight cultures of *P. ananatis* strains were spotted onto ABG agar plates. After incubation at 28°C for 24 h, the mucoid phenotype of colonies was investigated. For rugose phenotype, 5-μL overnight-cultured bacterial cells were inoculated onto LB agar plates and then incubated at 28°C for 48 h. The morphology of colonies was investigated.

### Swimming motility

The swimming motility test was performed in LB medium containing 0.3% Bacto agar. *P*. *ananatis* wild type, overexpression strains, and mutants were cultured in LB medium at 28°C for 24 h. The 1-mL overnight culture was harvested by centrifugation, washed twice with fresh LB medium, and resuspended in 100 µL of LB medium. An aliquot (1 µL) of each cell suspension was injected into the centers of swimming assay plates and incubated at 28°C for 24 h.

### Pellicle formation and degradation assays

Pellicle formation was carried out in LB medium. The overnight cultures of *P. ananatis* strains were diluted to an OD_600_ of 0.05 with fresh LB medium. Aliquots (3 mL) of diluted cultures were transferred to 24-well culture plates (Corning Inc., Corning, NY, USA) and incubated for 48 h at room temperature in static culture. The pellicle formation of cultures was investigated.

For pellicle degradation assays, pellicles were harvested from 2-day-old cultures and washed with phosphate-buffered saline followed by treatment with 0.86 U/mL of endo-1,4-β-D-glucanase (cellulase) or 0.1% (v/v) Proteinase K (Sigma-Aldrich, St. Louis, MO, United States) as described previously (38).

### Quantification of cellular c-di-GMP levels

The c-di-GMP was extracted as described previously with modification (71). *P. ananatis* PA13, overexpression strains, and mutants were cultured in LB medium at 28°C for 6 h to mid-exponential phase. At the indicated OD600 = 0.9, bacterial cultures were collected and washed with ice-cold phosphate-buffered saline (PBS) and pelleted. The pellets were resuspended with ice-cold PBS and incubated at 100°C for 5 min, and ice-cold ethanol to a final concentration of 65%. Samples were centrifuged, and the supernatant containing extracted c-di-GMP was transferred to a new microfuge tube. The cell pellets were used for the determination of total cellular protein levels. The supernatants were dried using a centrifugal evaporator and stored at -80°C until the next procedure. Samples resuspended with nanopure water were analyzed by high-performance liquid chromatography (Shimazu, Tokyo, Japan) equipped with an autosampler, degasser, pressure regulator, prefilter, and UV detector set to 254 nm. Separation was carried out using a YMC C18 column (250 × 4.6 mm; 5 µm) and a flow rate of 1mL/min. Commercially available c-di-GMP (Sigma-Aldrich) was used as a standard. The levels of cellular c-di-GMP were normalized to the corresponding total cellular protein levels.

### Virulence assays

Virulence of *P. ananatis* strains was evaluated in onion bulbs according to a previously described method (72). *P. ananatis* PA13, overexpression strains, and mutants were cultured in LB medium at 28°C overnight, centrifuged, washed, and suspended in sterilized distilled water (SDW) at OD_600_ = 0.8. Bacterial suspensions were infiltrated into onion bulbs using a syringe. SDW was used as a negative control. The inoculated onion bulbs were placed in a plastic box with saturated paper at room temperature. The disease symptoms of onion bulbs were observed at 15 days after inoculation.

## ACKNOWLEDGEMENTS

This research was supported by Basic Science Research Program through the National Research Foundation of Korea (NRF) funded by the Ministry of Education of the Republic of Korea (2015R1A6A1A03031413).

